# Sustained 1,25(OH)_2_D elevation promotes biphasic cholesterol elimination through bile acid and sterol pathways

**DOI:** 10.64898/2026.05.11.724189

**Authors:** Hirofumi Sogabe, Chiaki Abe, Eriko Takaramoto, Yo-ichi Nabeshima

**Affiliations:** Department of Aging Science and Medicine, Graduate School of Medicine, Kyoto University. 53 Shogoin Kawahara-cho, Sakyo-ku, Kyoto 606-8507, JAPAN

**Author notes:** These authors contributed equally to this study (Hirofumi Sogabe, Chiaki Abe).

## Abstract

Lower circulating levels of 25-hydroxyvitamin D (25(OH)D) have been associated with adverse lipid profiles in humans; however, the underlying mechanisms remain poorly defined. Here, we investigated how sustained elevation of active vitamin D, 1,25-dihydroxyvitamin D (1,25(OH)₂D), promotes cholesterol elimination using two complementary mouse models: *α-Klotho* KO mice, which exhibit chronically elevated endogenous 1,25(OH)₂D, and WT mice repeatedly administered 1,25(OH)₂D₃. *α-Klotho* KO mice showed increased fecal elimination of cholesterol and total bile acids, accompanied by increased expression of hepatic *Sr-b1*, *Abcg5/Abcg8*, and *Cyp7a1* and reduced expression of duodenal *Npc1l1* and ileal *Asbt*. The 1,25(OH)₂D₃-administered model revealed a biphasic cholesterol elimination response involving an early bile acid route and a later neutral sterol route. In the early phase (day 3), fecal elimination of bile acids increased, coinciding with reduced hepatic bile acid levels and increased hepatic *Mrp2* expression, whereas ileal *Asbt*/ASBT expression and early induction of genes related to bile acid synthesis were not observed. In the later phase (days 12–15), fecal cholesterol elimination increased, accompanied by induction of hepatic cholesterol uptake and biliary secretion-related pathways, including increased *Sr-b1* and *Abcg5/Abcg8* expression and elevated SR-BI protein levels, together with reduced duodenal *Npc1l1* expression. *Abca1* was significantly induced in skeletal muscle, suggesting additional regulation of peripheral cholesterol metabolism. These findings show that sustained elevation of 1,25(OH)₂D promotes fecal elimination of cholesterol through coordinated hepatic, intestinal, and peripheral regulation. These results provide a mechanistic framework for understanding the influence of active vitamin D on systemic cholesterol homeostasis.

## INTRODUCTION

Cholesterol homeostasis is maintained through coordinated regulation of its intestinal absorption, hepatic uptake, *de novo* synthesis, lipoprotein-mediated transport, and elimination via hepatobiliary and intestinal routes (1–3). Perturbation of this balance is implicated in cardiometabolic disorders, including atherosclerosis and nonalcoholic fatty liver disease (NAFLD) (1,4). Because mammals lack a major pathway to degrade the sterol ring structure, excess cholesterol must be eliminated primarily by (i) direct secretion of cholesterol into bile, followed by elimination of cholesterol in feces, and (ii) conversion of cholesterol into bile acids, followed by fecal elimination of bile acids (1,5). At the molecular level, secretion of hepatobiliary cholesterol involves hepatic uptake of circulating lipoprotein cholesterol and canalicular export into bile. Scavenger receptor class B type 1 (SR-BI) contributes to hepatic uptake of HDL-derived cholesteryl esters (6,7), whereas the ABCG5/ABCG8 heterodimer promotes excretion of hepatobiliary cholesterol and increases fecal elimination of cholesterol (8). Cholesterol is converted into bile acids by cholesterol 7α-hydroxylase (CYP7A1), a rate-limiting enzyme in the classical bile acid synthesis pathway, which is regulated by bile acid–nuclear receptor feedback mechanisms (9,10). Bile acids are secreted into bile mainly via the bile salt export pump (BSEP) and are efficiently reabsorbed in the ileum by the apical sodium-dependent bile acid transporter (ASBT) to maintain enterohepatic circulation (11). In the small intestine, luminal absorption of food-derived and biliary-secreted cholesterol is mediated by the NPC1-like intracellular cholesterol transporter 1 (NPC1L1) (12).

The *α-Klotho* KO mouse, characterized by its short lifespan, exhibits numerous histological and serological phenotypes associated with human aging. These phenotypes include calcification of various soft tissues, atherosclerosis, bone malformation, pulmonary emphysema, kyphosis, senile skin atrophy, severe reduction in white fat deposits, and infertility (13,14). *α-Klotho* KO mice appear almost normal until 3 weeks of age, after which they display growth retardation; their body weights remain under 10 g throughout their life spans under standard nutritional and exercise conditions, and they die prematurely (average life span of 70–80 days). Serum levels of 1,25-dihydroxyvitamin D (1,25(OH)_2_D) in *α-Klotho* KO mice are much higher than those in WT mice at 2 weeks of age before the occurrence of multiple histological phenotypic changes and remain significantly higher throughout their lifespan (14). Such high serum levels of 1,25(OH)_2_D are closely associated with multiple human aging-related pathological phenotypes and the dysregulation of serum levels of calcium, phosphate, fibroblast growth factor 23 (FGF23), glucose, and triglycerides (TGs) (15). Additionally, reducing serum levels of 1,25(OH)_2_D in *α-Klotho* KO mice, either by restriction of vitamin D (vitamin D-deficient diet and housing mice under shaded conditions) or by genetic manipulation (crossing *α-Klotho* KO mice with *cyp27b1* KO mice), can improve many of the pathological and serological phenotypes (15,16), suggesting that chronically elevated 1,25(OH)_2_D is the key determinant of *α-Klotho*-deficient phenotypes. Therefore, *α-Klotho* KO provides a useful model for studying long-term effects of elevated signaling of active vitamin D.

Vitamin D plays a central role in the regulation of mineral homeostasis, particularly by maintaining the balance of calcium and phosphate (17). However, increasing evidence indicates that vitamin D and the vitamin D receptor (VDR) are involved in many physiological processes beyond mineral homeostasis (18,19). Notably, vitamin D deficiency is associated with increased levels of circulating total cholesterol and TGs (20–22), and vitamin D supplementation may improve lipid profiles in vitamin D-deficient individuals (23). The importance of 1,25(OH)_2_D in regulation of cholesterol has also been reported in mice. Crossing *α-Klotho* KO mice with ob/ob mice significantly ameliorated deposition of cholesterol in the liver; however, it did not ameliorate increased blood cholesterol levels, although both are typical phenotypes observed in ob/ob mice (24). This observation suggests that elevated levels of 1,25(OH)₂D regulate hepatic cholesterol metabolism by enhancing cholesterol disposal and/or suppressing synthesis of *de novo* cholesterol. Together, these observations imply that the status of vitamin D is linked to systemic cholesterol metabolism; however, the mechanism of action remains poorly understood. To the best of our knowledge, only limited mechanistic insights have been reported, including the effects of vitamin D on hepatic CYP7A1 (25,26). Based on the above observations, we investigated the comprehensive actions of 1,25(OH)_2_D in the regulation of cholesterol metabolism using two complementary approaches: a genetic model with chronic endogenous elevation of 1,25(OH)_2_D (*α-Klotho* KO mice) and a pharmacological model related to administration of 1,25-dihydroxyvitamin D_3_ (1,25(OH)_2_D_3_) in C57BL/6J mice. In particular, we focused on (i) hepatic cholesterol uptake- and export-related gene expression, (ii) pathway related to *de novo* cholesterol synthesis, (iii) intestinal cholesterol absorption- and bile acid reabsorption-related gene expression, and (iv) temporal changes that lead to enhanced elimination of cholesterol.

Here, we propose that sustained elevation of 1,25(OH)_2_D_3_ levels promotes cholesterol elimination through an early increase in fecal loss of bile acids, followed by a subsequent increase in fecal loss of neutral sterols, thus providing a mechanistic framework for interpreting the mechanism by which 1,25(OH)_2_D affects cholesterol metabolism.

## Materials and Methods

### Animal Preparation

*α-klotho* KO mice were generated by crossing heterozygous *α-klotho* mice. Genotyping of *α-klotho* mice was performed using tail biopsies collected at three weeks of age. DNA was amplified following a three-step cycle protocol with KOD FX polymerase using the primers listed in Supplementary Table 1. Male *α-klotho* KO and wild-type mice aged 6–8 weeks were used in the experiments. For experiments involving calcitriol administration, 5-week-old C57BL/6J mice were purchased from Japan SLC (Shizuoka, Japan) and acclimatized for one week prior to use. Calcitriol (Rocaltrol®; Kyowa Kirin) was administered intraperitoneally every 12 h. The mice were anesthetized, and blood was collected via cardiac puncture using heparinized syringes, followed by perfusion with cold phosphate-buffered saline (PBS). The liver and small intestine were harvested, and the small intestine was divided into three equal segments: the duodenum (closest to the stomach), jejunum, and ileum. The organs were snap-frozen in liquid nitrogen and stored at − 80 °C. Feces from singly housed mice were collected every 24 h and stored at – 80 °C until use. All mice were housed under a 12-hour light/dark cycle with ad libitum access to food and water. Experiments were conducted using freely fed male mice under protocols approved by the Animal Care and Use Committee of the Kyoto University Graduate School of Medicine in accordance with institutional regulations and guidelines.

### Total Cholesterol Measurement

The feces were ground into powder using a mortar and pestle. Liver and fecal samples were homogenized in a solution of chloroform: isopropanol:NP-40 (7:11:0.1, v/v/v) using a microhomogenizer. The homogenates were centrifuged at 15,000 × g for 10 min, and the supernatants were collected. Lipids were extracted by vacuum centrifugation at 50 °C and 3,000 × g for 30 min. The resulting lipids were resuspended in 1× assay diluent from the Total Cholesterol Assay Kit (colorimetric method; Cell BioLabs, STA-384), and the total cholesterol concentration was measured according to the manufacturer’s protocol.

### Total Bile Acid Measurement

Feces were collected and prepared as described above for cholesterol measurement. Liver and fecal samples were homogenized in isopropanol using a microhomogenizer. After centrifugation at 15,000 × g for 10 min, the organic phase was diluted, and the total bile acid concentration was measured using a Total Bile Acid Assay Kit (Cell BioLabs, STA-631), following the manufacturer’s instructions.

### RNA Extraction and qPCR Analysis

Total mRNA was extracted from the tissues or cells using the RNeasy® Plus Mini Kit (QIAGEN, 74134) according to the manufacturer’s protocol. Complementary DNA (cDNA) was synthesized from mRNA using the ReverTra Ace® qPCR RT Master Mix (TOYOBO, FSQ-201). Quantitative PCR (qPCR) was performed in duplicate for each sample using the Step One Plus System (Applied Biosystems, USA) and THUNDERBIRD® SYBR™ qPCR Mix (TOYOBO, QPS-201), with the primers listed in Supplementary Table 2. Gene expression levels were normalized to β-actin, and relative mRNA expression was calculated using the 2^−ΔΔCt method.

### Biochemical analysis

Heparinized blood was centrifuged at 3,000 x g for 15 min at 4℃, and the supernatant was collected. Plasma cholesterol and HDL-cholesterol concentration were measured using a DRI-CHEM 3000 (FUJIFILM).

### Western blot

The tissues were lysed in RIPA buffer containing protease inhibitors. Homogenates were incubated on ice for 30 min, then centrifuged at 16,000 x g for 10 min at 4℃. The supernatants were collected and protein concentrations were determined using the Pierce™ BCA Protein Assay Kit (Pierce, Cat No. 23227). The protein was mixed with 6 × SDS sample buffer (Final conc; 0.35 M Tris-HCl, 10% SDS, 30% glycerol, 9.3% DTT, 0.012% BPB), and boiled at 95℃ for 5 min. Denatured protein was subjected to SDS-PAGE and transferred to a polyvinylidene fluoride (PVDF) membrane. Membranes were washed with TBS-T for 5 min, then blocked with 5% skim milk in TBS-T for 1 h at r.t. The membranes were washed with TBS-T (3 times) for 10 min and incubated with anti-SR-BI (Abcam, ab52629, 1:1,000), anti-ABCG5 (Protein tech, 27722-1-AP, 1:1,000), anti-ABCG8 (Novus, NBP1-71706, 1:1,000), anti-ASBT (Protein tech, 25245-1-AP,1:1,000), or anti-β-actin (CST, 4967,1:1,000) for overnight at 4℃. Membranes were washed with TBS-T (3 times) and incubated with anti-rabbit HRP conjugated secondary antibody (Cytiva, NA934V, 1:10,000) and anti-mouse HRP conjugated secondary antibody (Cytiva, NA9310V, 1:10,000) in 5% skim milk in TBS-T for 1 h at r.t. Membranes were washed with TBS-T (3 times), then incubated with ECL™ Prime Western Blotting Detection Reagent (Cytiva, RPN2236) for 5 min and detected with LAS500 (Cytiva).

### Statistical Analysis

The data are presented as the mean ± standard deviation (SD). Potential outliers were identified using Dixon’s Q-test and excluded when necessary. The equality of variances was evaluated using the F-test before conducting the t-test. Student’s t-test was applied when variances were equal, while Welch’s t-test was employed for unequal variances. Statistical significance was determined at P < 0.05.

## RESULTS

### Enhanced fecal elimination of cholesterol and bile acids in *α-Klotho* KO mice

A previous study showed that accumulation of hepatic cholesterol in ob/ob mice was markedly reduced when these mice were crossed with *α-Klotho* KO mice (24), which exhibited chronically elevated circulating levels of 1,25(OH)₂D. This finding suggests that elevated 1,25(OH)₂D levels lower hepatic cholesterol levels through one or more of the following mechanisms: (i) enhanced elimination of cholesterol and bile acids in feces, (ii) suppression of *de novo* cholesterol synthesis, and/or (iii) coordinated changes in hepatic cholesterol uptake and export. Therefore, we first examined fecal elimination of cholesterol and total bile acids. Compared with WT mice, *α-Klotho* KO mice showed a significant increase in the fecal elimination of cholesterol and bile acids (Fig. 1A–B). These findings indicate that chronic elevation of 1,25(OH)₂D is associated with enhanced fecal elimination of cholesterol and bile acids.

**Fig. 1.**
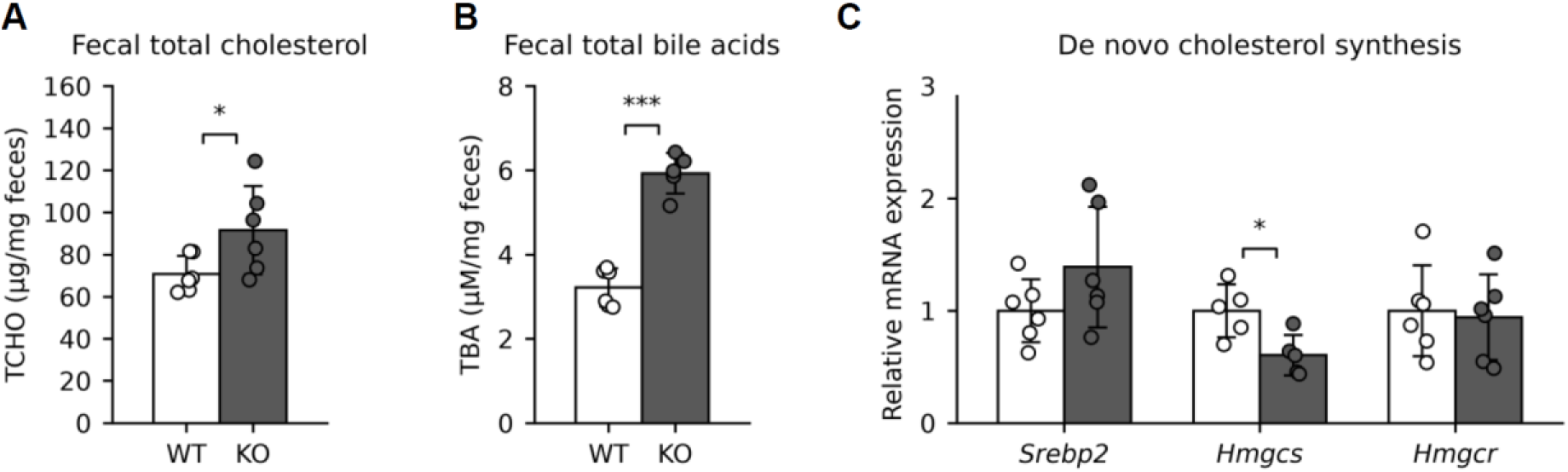
Cholesterol and bile acid contents and expression of genes involved in *de novo* cholesterol synthesis in *α-Klotho* KO mice. Fecal samples were collected over 24 h, and the total bile acid and cholesterol contents in the feces were measured (A–B). mRNA expression in the *de novo* cholesterol synthesis pathway (C). The white and dark gray bars indicate WT and *α-Klotho* KO mice, respectively. The mRNA expression levels were normalized to β-actin. Values are presented as mean ± SD (n = 5–6); * p < 0.05, *** p < 0.001.

Increased fecal elimination of cholesterol may be accompanied by compensatory changes in synthesis of hepatic cholesterol (27,28). Thus, we examined whether genes related to biosynthesis of hepatic *de novo* cholesterol were transcriptionally upregulated in *α-Klotho* KO mice. No significant increases were observed in the hepatic expression levels of *Srebp2*, a master regulator of *de novo* cholesterol biosynthesis, *HMG-CoA reductase* (*Hmgcr*), the rate-limiting enzyme in the cholesterol biosynthesis pathway, or *HMG-CoA synthase* (*Hmgcs*), which is involved in cholesterol synthesis (Fig. 1C). These findings indicate that the increased fecal elimination of cholesterol and bile acids in *α-Klotho* KO mice is unlikely to be explained by transcriptional upregulation of hepatic *de novo* cholesterol biosynthesis.

### *α-Klotho* deficiency alters hepatic cholesterol transport and bile acid synthesis pathways

Fecal elimination of cholesterol and bile acids can be increased not only by enhanced cholesterol synthesis but also by the alterations in hepatic cholesterol transport, biliary cholesterol secretion, and/or conversion of cholesterol into bile acids. Therefore, we next examined the expression of hepatic genes involved in these pathways.

Expression of hepatic *Sr-b1* was significantly increased in *α-Klotho* KO mice (Fig. 2A). Additionally, expression of hepatic *Abcg5* and *Abcg8*, which promote biliary cholesterol secretion, was also significantly increased in *α-Klotho* KO mice (Fig. 2A). These results indicate that hepatic cholesterol transport is upregulated in *α-Klotho* KO mice. Next, we examined the pathways of bile acid synthesis. Expression of hepatic *Cyp7a1* was significantly increased in *α-Klotho* KO mice, whereas that of *sterol 26-hydroxylase* (*Cyp27a1*), which is involved in an alternative bile acid synthesis pathway, remained unchanged (Fig. 2B). Expression of *Bsep*, which mediates canalicular bile acid secretion, was not altered significantly, whereas that of *ATP-binding cassette sub-family C member 2* (*Mrp2*), a canalicular organic anion transporter, tended to increase (p = 0.057) (Fig. 2C). Furthermore, the expression of *Mrp3* and *Mrp4*, the basolateral efflux transporters, was significantly increased (Fig. 2C). These results indicate that pathways involved in hepatic bile acid synthesis and transport are upregulated in *α-Klotho* KO mice. Among the other transporters and receptors, expression of hepatic *Abca1* was increased, whereas that of LDL receptor (*Ldlr*) and hepatic sodium/bile acid cotransporter (*Ntcp*) remained unchanged (Fig. 2A and C).

**Fig. 2.**
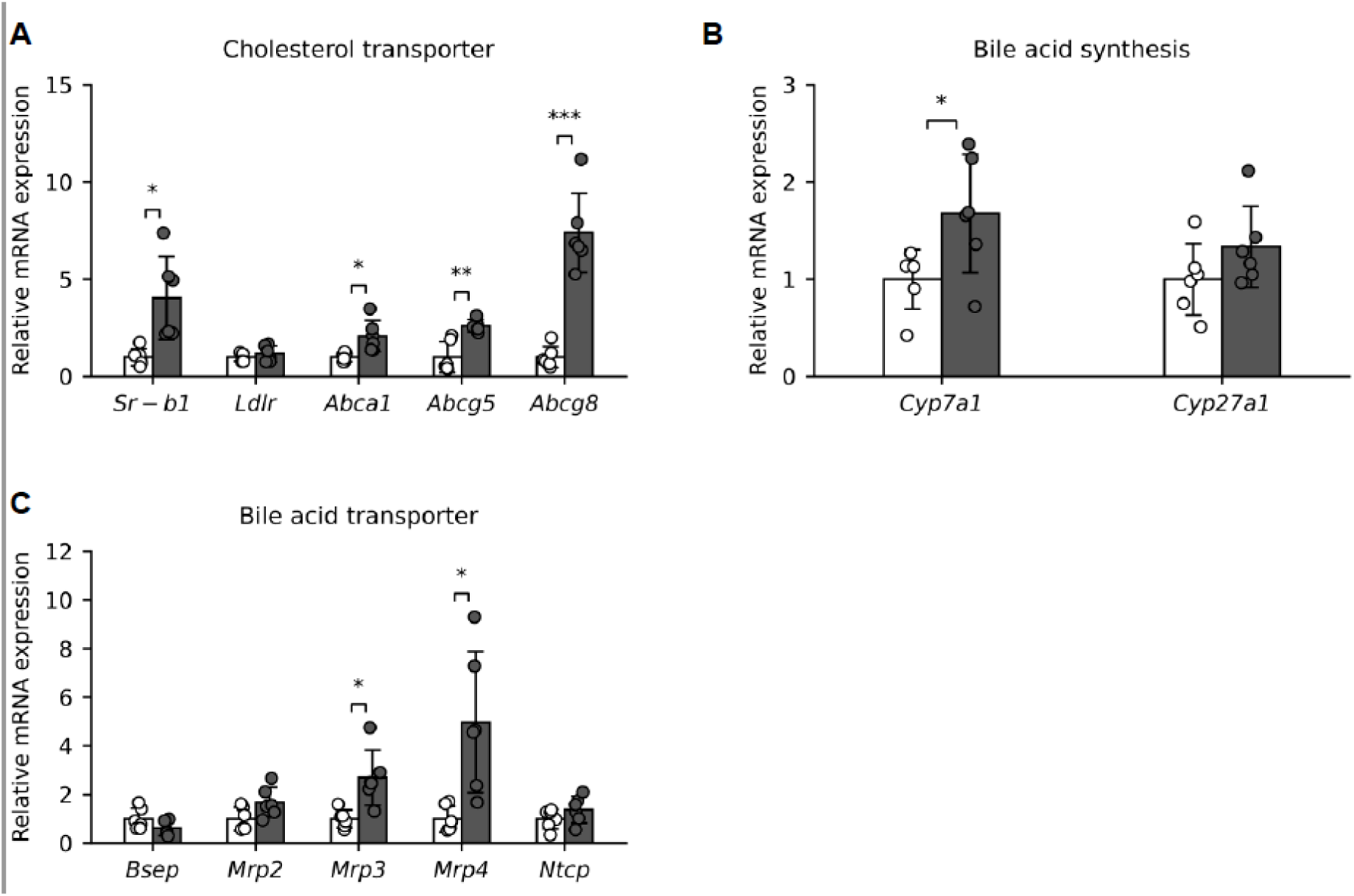
mRNA expression in the livers of *α-Klotho* KO mice. Cholesterol influx and efflux transporters (A), bile acid synthesis (B), and bile acid influx and efflux transporters (C). The white and dark gray bars indicate WT and *α-Klotho* KO mice, respectively. mRNA expression levels were normalized to β-actin and are presented as mean ± SD (n = 5–6); * p < 0.05, ** p < 0.01, *** p < 0.001.

Taken together, these results indicate that *α-Klotho* KO mice exhibit coordinated transcriptional changes in hepatic cholesterol and bile acid metabolism. In particular, the increased expression of *Sr-b1*, *Abcg5*, *Abcg8*, and *Cyp7a1* is consistent with the transcriptional remodeling of pathways involved in hepatic uptake of HDL-derived cholesterol, biliary cholesterol secretion, and classical bile acid synthesis.

### Suppressed intestinal absorption/reabsorption is associated with increased fecal cholesterol elimination in ***α****-Klotho* KO mice

To determine whether changes in intestinal absorption of cholesterol and bile acids contribute to the enhanced elimination of cholesterol and bile acids, we examined the expression levels of *Npc1l1*, a cholesterol uptake transporter, and *Asbt*, a bile acid uptake transporter. Expression levels of *Npc1l1* in the duodenum and jejunum and *Asbt* in the duodenum and ileum were significantly reduced in *α-Klotho* KO mice (Fig. 3A–B). Together, these results imply that enhanced fecal elimination of cholesterol and bile acids in *α-Klotho* KO mice is associated with coordinated transcriptional changes that promote disposal of hepatic cholesterol and bile acids while suppressing their absorption and reabsorption in the small intestine.

**Fig. 3.**
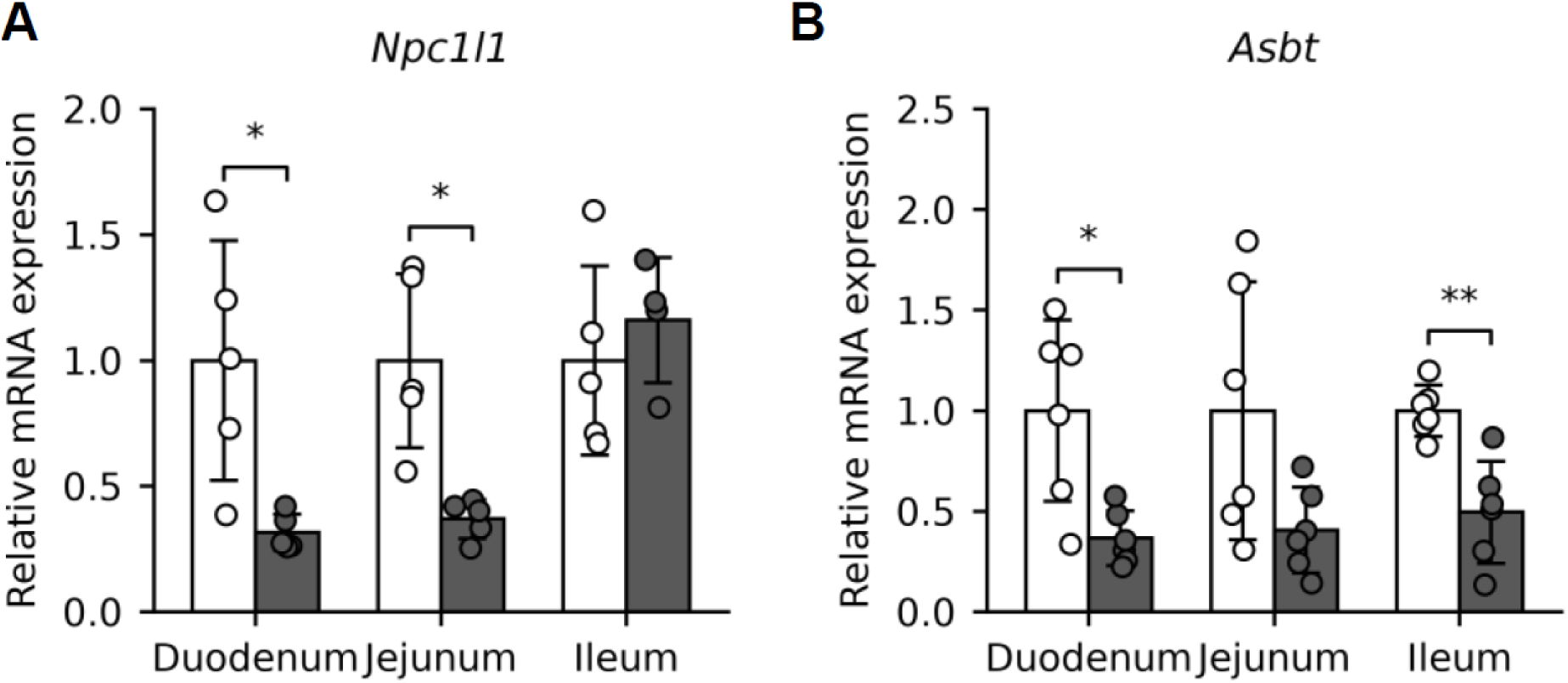
mRNA expression in the small intestine of *α-Klotho* KO mice. Cholesterol influx transporters and bile acid influx transporters in the small intestine (A–B). The white and dark gray bars indicate WT and *α-Klotho* KO mice, respectively. mRNA expression levels were normalized to β-actin and are presented as mean ± SD (n = 5–6); * p < 0.05, ** p < 0.01.

### 1,25(OH)2D3 promotes early-phase fecal elimination of bile acids

Although *α-Klotho* KO mice have chronically elevated circulating levels of 1,25(OH)₂D, *α-Klotho* deficiency itself or other secondary factors also contribute to changes in cholesterol and bile acid metabolism. To test whether elevated 1,25(OH)_2_D_3_ levels were sufficient to induce the fecal elimination phenotype, C57BL/6J mice were administered 1,25(OH)_2_D_3_ (twice/day) and analyzed on days 1, 2, 3, 6, 9, 12, and 15 (Fig. 4A). Consistent with findings from *α-Klotho* KO mice, fecal bile acid levels significantly increased on day 3 and remained elevated at subsequent time points (Fig. 4B). Notably, hepatic bile acid levels showed an overall decreasing trend from day 3 (Fig. 4C), and the decreased bile acid levels in the liver and increased fecal elimination of bile acids were consistent with these findings.

**Fig. 4.**
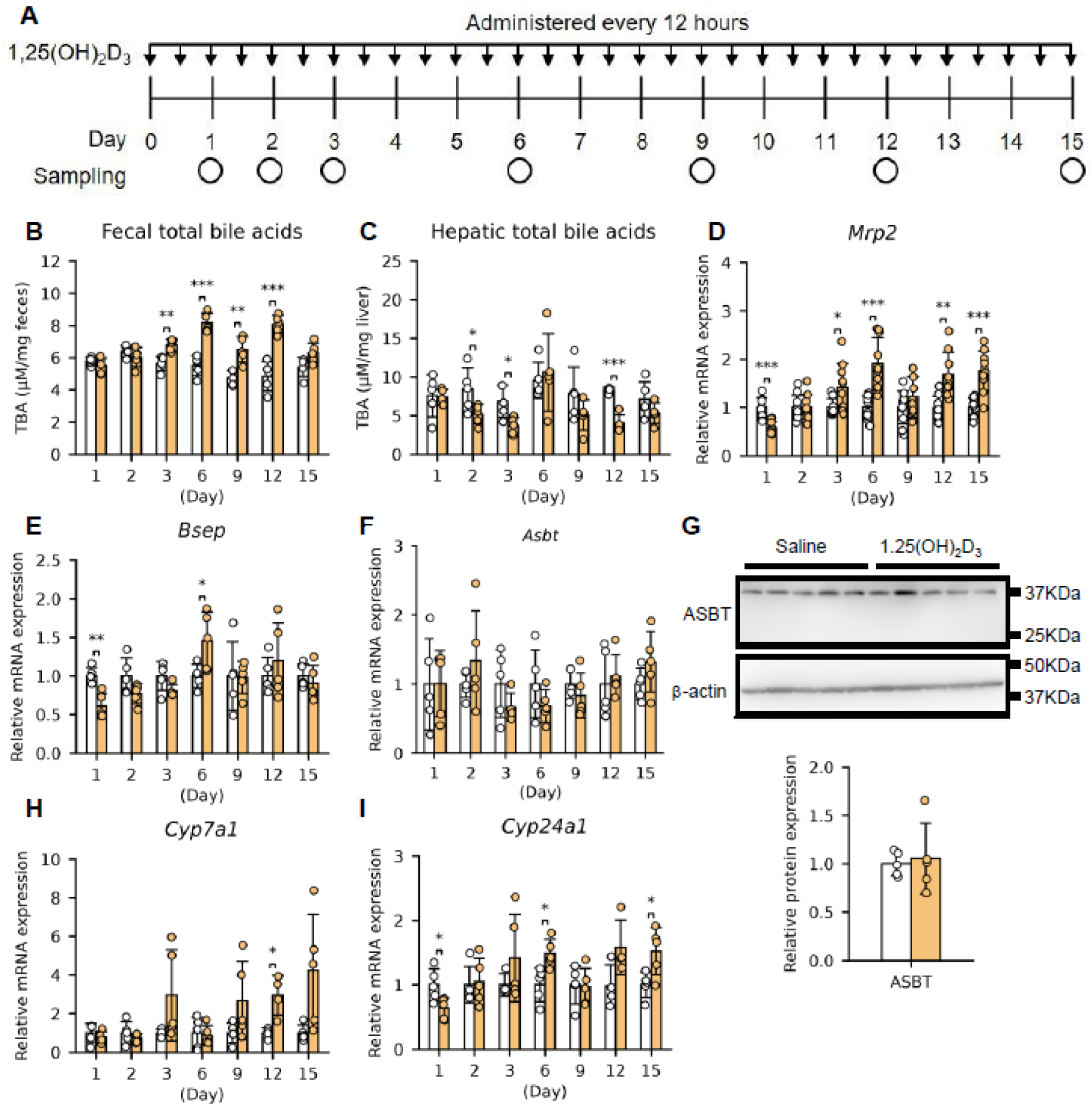
Effect of 1,25(OH)_2_D_3_ on bile acid metabolism. 1,25(OH)_2_D_3_ administration regimen (A), total bile acids in the feces and liver (B–C), mRNA expression of hepatic bile acid efflux transporters (D–E), mRNA and protein expression of ileal bile acid influx transporters (F–G), and mRNA expression of bile acid synthesis (H–I). The white and orange bars indicate saline and 1,25(OH)_2_D_3_ administered mice, respectively. The mRNA and protein expression levels were normalized to β-actin. Values are presented as mean ± SD (n = 4–5); * p < 0.05, ** p < 0.01, *** p < 0.001.

In the liver, expression of *Mrp2* increased from day 3 and continued to increase at subsequent time points, whereas consistent increases in *Bsep* expression were not observed (Fig. 4D–E). Ileal *Asbt* expression levels were not significantly altered at any time point (Fig. 4F). Furthermore, ASBT protein levels were not altered on day 3, at which fecal bile acid elimination had already increased (Fig. 4G). These findings suggest that an early increase in fecal bile acid elimination is associated with *Mrp2* induction and is unlikely to be explained by suppression of ileal bile acid reabsorption. *Cyp7a1* and *Cyp27a1* expression was induced at later time points, mainly on days 12–15 (Fig. 4H– I). The delayed induction of *Cyp7a1* and *Cyp27a1* further suggests that increased bile acid synthesis is unlikely to be the primary driver of the early-phase increase in fecal bile acid elimination. Collectively, these results support that an early increase in fecal bile acid elimination following 1,25(OH)₂D₃ administration is more closely associated with disposal of hepatic bile acids rather than with reduced bile acid reabsorption in the intestine or early induction of bile acid synthesis.

### 1,25(OH)₂D₃ administration promotes late-phase fecal cholesterol elimination and coordinated regulation of cholesterol metabolism

Following the early increase in fecal bile acid elimination, we examined whether 1,25(OH)₂D₃ administration altered cholesterol metabolism. Fecal total cholesterol levels increased significantly on day 15 (Fig. 5A). In contrast, the hepatic total cholesterol content began to decline from day 6 and was significantly reduced at later time points (Fig. 5B). Thus, the reduction in hepatic cholesterol content occurred after the increase in fecal bile acid elimination and preceded the significant increase in fecal cholesterol elimination. Because administration of 1,25(OH)₂D₃ induced temporal changes in hepatic cholesterol content, we examined the expression of hepatic genes involved in cholesterol uptake and biliary cholesterol secretion. Expression of hepatic *Sr-b1* increased from day 6, and that of *Abcg5* and *Abcg8* also increased or tended to increase during the later phases (Fig. 5C–E). On day 15, hepatic SR-BI protein levels increased (Fig. 5F). Additionally, ABCG5 and ABCG8 protein levels tended to increase, although the extent of induction varied across experiments. (Fig.5F and Supplementary Fig. 2A–C). These findings support the late-phase remodeling of the hepatic pathways involved in HDL-derived cholesterol uptake and biliary cholesterol secretion.

**Fig. 5.**
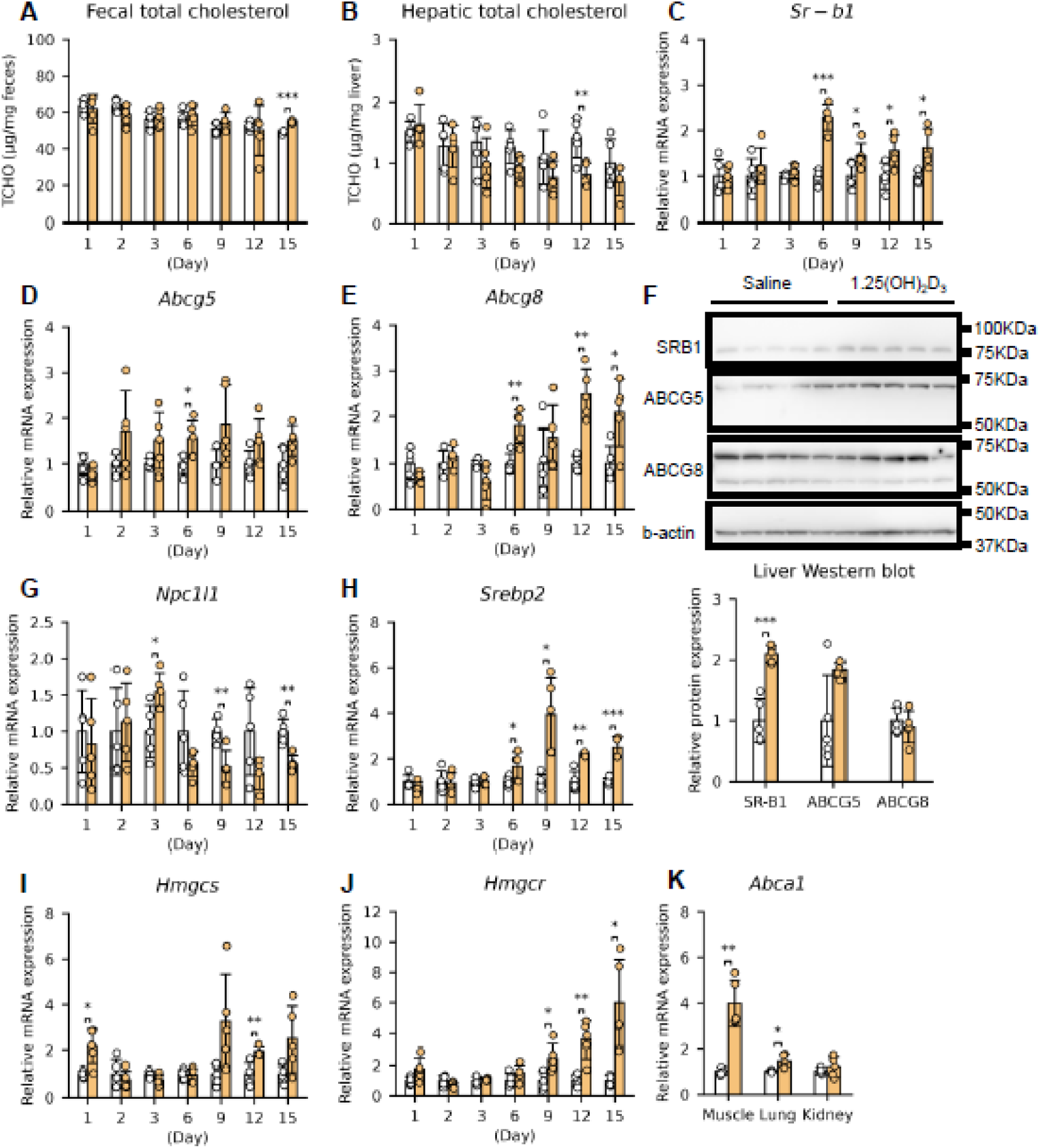
Effect of 1,25(OH)_2_D_3_ on cholesterol metabolism. Total cholesterol levels in the feces and liver (A–B). mRNA and protein expression of cholesterol efflux and influx transporters in the liver (C–F). mRNA expression of cholesterol influx transporter in the duodenum (G). mRNA expression of de novo cholesterol synthesis in the liver (H–J). mRNA expression of cholesterol efflux transporters in peripheral tissues (K). The white and orange bars indicate saline and 1,25(OH)_2_D_3_ administered mice, respectively. The mRNA and protein expression levels were normalized to β-actin. Values are presented as mean ± SD (n = 4–5); * p < 0.05, ** p < 0.01, *** p < 0.001.

Because intestinal cholesterol absorption is mediated primarily by duodenal NPC1L1, we assessed *Npc1l1* expression. Expression of duodenal *Npc1l1* began to decrease from day 6 and was significantly reduced at multiple time points (Fig. 5G). These findings suggest that administration of 1,25(OH)₂D₃ reduces intestinal cholesterol absorption concurrently with an increase in fecal cholesterol elimination.

We then examined whether hepatic *de novo* cholesterol biosynthesis was altered in response to reduction in hepatic cholesterol content and enhanced fecal cholesterol elimination. Hepatic expression of *Srebp2* began to increase on day 6, subsequently leading to induction of *Hmgcs* and *Hmgcr*, which was predominantly observed between days 9 and 15 (Fig. 5H–J). Because these genes were induced after hepatic cholesterol content began to decline, their upregulation implied a compensatory response to hepatic cholesterol depletion and enhanced cholesterol disposal.

We also examined whether administration of 1,25(OH)₂D₃ affected cholesterol metabolism in peripheral tissues such as the quadriceps muscle, lungs, and kidneys on day 15. Expression of *Abca1* was markedly increased in the quadriceps muscle and modestly increased in the lungs, whereas no apparent induction was observed in the kidneys (Fig. 5K). Serum total cholesterol and HDL-C levels were largely unchanged (Supplementary Fig.2A–B). These findings suggest that the administration of 1,25(OH)₂D₃ exerts broader effects on cholesterol metabolism, including in peripheral tissues, without significantly altering circulating total cholesterol or HDL-C levels.

Collectively, these results show that 1,25(OH)₂D₃ administration induces late-phase fecal cholesterol elimination, accompanied by the coordinated regulation of hepatic, intestinal, and peripheral cholesterol metabolism. Together with the preceding increase in fecal bile acid elimination, these findings provide a model in which 1,25(OH)₂D₃ first promotes cholesterol disposal through the bile acid pathway and subsequently enhances fecal cholesterol elimination through hepatic–intestinal remodeling with additional regulation of peripheral cholesterol metabolism.

## Discussion

In this study, we demonstrated that sustained elevation of circulating 1,25(OH)_2_D levels promotes fecal elimination of cholesterol and bile acids using two complementary models: (i) *α-Klotho* KO mice with chronically elevated endogenous 1,25(OH)_2_D and (ii) WT mice administered 1,25(OH)_2_D_3_. In *α-Klotho* KO mice, fecal elimination of both cholesterol and bile acids was increased and accompanied by coordinated transcriptional changes in the hepatic and intestinal pathways involved in cholesterol and bile acid metabolism. The mice administered 1,25(OH)_2_D_3_ recapitulated the key features of the chronic exposure phenotype and enabled the clarification of temporal changes in the metabolic response.

A major finding of this study is that administration of 1,25(OH)₂D₃ induced a two-phase response in fecal elimination of bile acids and cholesterol: (i) fecal elimination of bile acids increased by day 3 (early-phase response) and (ii) fecal elimination of cholesterol increased by day 15 (late-phase response). Hepatic cholesterol content began to decline between these two events, after an increase in fecal elimination of bile acids and before a significant increase in fecal elimination of cholesterol. This temporal pattern reveals that 1,25(OH)₂D₃ first promotes cholesterol disposal through the bile acid pathway and subsequently enhances fecal cholesterol elimination through coordinated transcriptional regulation of genes involved in hepatic and intestinal cholesterol metabolism.

The early increase in fecal bile acid elimination was associated with induction of hepatic *Mrp2*, while a sustained induction of *Bsep* was not observed. Although BSEP is the principal canalicular bile acid exporter, the absence of sustained *Bsep* induction implies the involvement of different mechanisms related to the control of early fecal bile acid elimination. MRP2 is involved in bile formation, particularly through glutathione-dependent bile acid-independent bile flow; diminished MRP2 function is associated with impaired bile flow and decreased biliary glutathione excretion (29). Furthermore, *Mrp2*-deficient rats demonstrate impaired biliary bile acid secretion, indicating that MRP2 may affect hepatic bile acid disposal (30). Consequently, the induction of *Mrp2* observed in the 1,25(OH)₂D₃ administration model may enhance bile formation and increase the excretion of bile acid-containing bile into the intestinal lumen, thereby promoting fecal bile acid elimination. Conversely, *Asbt* gene and ASBT protein expression were not altered in the 1,25(OH)₂D₃ administration model, and ileal *Asbt* expression was reduced rather than increased in α-Klotho KO mice. Thus, suppression of ASBT-dependent ileal bile acid reabsorption is unlikely to explain the early increase in fecal bile acid elimination. These observations support that enhanced hepatic bile acid hepatobiliary excretion, rather than suppression of ileal bile acid reabsorption, predominantly contributes to the early increase in fecal bile acid elimination following 1,25(OH)₂D₃ administration.

The late-phase increase in fecal cholesterol elimination was accompanied by increased hepatic *Sr-b1* gene and SR-BI protein expression, increased *Abcg5* and *Abcg8* gene expression, an increasing trend in ABCG5 and ABCG8 protein levels, and reduced duodenal *Npc1l1* expression. Collectively, these findings are consistent with increased hepatic cholesterol uptake/secretion capacity and reduced intestinal cholesterol absorption. Furthermore, the late induction of *Srebp2*, *Hmgcs*, and *Hmgc*r was consistent with the compensatory activation of *de novo* cholesterol biosynthesis in response to hepatic cholesterol depletion and enhanced cholesterol elimination. In addition to hepatic and intestinal responses, administration of 1,25(OH)₂D₃ markedly induced *Abca1* expression in the quadriceps muscle and modestly increased *Abca1* expression in the lungs, suggesting that elevated 1,25(OH)₂D₃ may affect not only the liver and intestines but also peripheral cholesterol metabolism.

Our findings also provide a mechanistic framework for interpreting observations in human cohort studies regarding the relationship between blood 25(OH)D and cholesterol levels. Observational studies have reported higher cholesterol and TG levels in individuals with vitamin D deficiency or lower serum 25(OH)D levels, and interventional studies have indicated that supplementation with vitamin D or 25(OH)D can decrease blood cholesterol levels in vitamin D-deficient individuals. However, mechanisms underlying this cholesterol-lowering effect remain unclear. The present study suggests that sustained elevation of 1,25(OH)₂D₃ can shift cholesterol metabolism from the basal homeostatic state toward an enhanced fecal elimination state through coordinated regulation of hepatic, intestinal, and peripheral pathways. This includes early enhancement of fecal bile acid elimination, late-phase induction of hepatic cholesterol uptake and biliary secretion-related pathways, suppression of intestinal cholesterol absorption, and induction of peripheral cholesterol secretion.

The development of cholesterol-lowering drugs has targeted specific components of cholesterol metabolism, such as statins, ezetimibe, and bile acid sequestrants (32–34). In contrast, 1,25(OH)_2_D_3_ coordinately regulates multiple components of cholesterol and bile acid metabolism. However, owing to the side effects of 1,25(OH)_2_D_3_, particularly hypercalcemia, its use in cholesterol therapy poses certain risks to individuals. Therefore, identification and targeting of downstream pathways that preserve the cholesterol elimination-enhancing effects of 1,25(OH)₂D₃ while avoiding calcemic toxicity may be a promising strategy for cholesterol regulation.

This study has several limitations. (i) Our mechanistic interpretation is based primarily on transcript measurements, analyses of selected proteins, and measurements of cholesterol and total bile acids in the liver and feces. Direct measurements of biliary cholesterol and bile acid levels were not performed. (ii) Although ASBT protein expression was examined, protein levels of all transporters, including MRP2 and NPC1L1, were not fully assessed. (iii) The possible involvement of peripheral cholesterol mobilization was inferred from gene expression changes, including *Abca1* induction; however, this was not directly tested using tracer-based cholesterol flux assays.

In conclusion, sustained elevation of 1,25(OH)₂D in *α-Klotho* KO mice promotes fecal elimination of bile acids and cholesterol through temporally distinct and coordinated regulation of cholesterol metabolism (Fig. 1A, B). In C57BL/6J mice, repeated 1,25(OH)₂D₃ administration induced sequential metabolic events, including (i) elimination of bile acids to feces (Fig. 4B), (ii) reduction of hepatic cholesterol (Fig. 5B), (iii) suppression of intestinal cholesterol absorption (Fig. 5G), and (iv) elimination of fecal cholesterol to feces (Fig. 5A). Together, these observations indicate that 1,25(OH)₂D₃ is involved in the broader changes in hepatic and peripheral cholesterol metabolism (Fig. 6). These findings expand the physiological significance of vitamin D signaling beyond mineral homeostasis and suggest that vitamin D acts as an endocrine regulator that coordinates lipid utilization and sterol elimination.

**Fig. 6.**
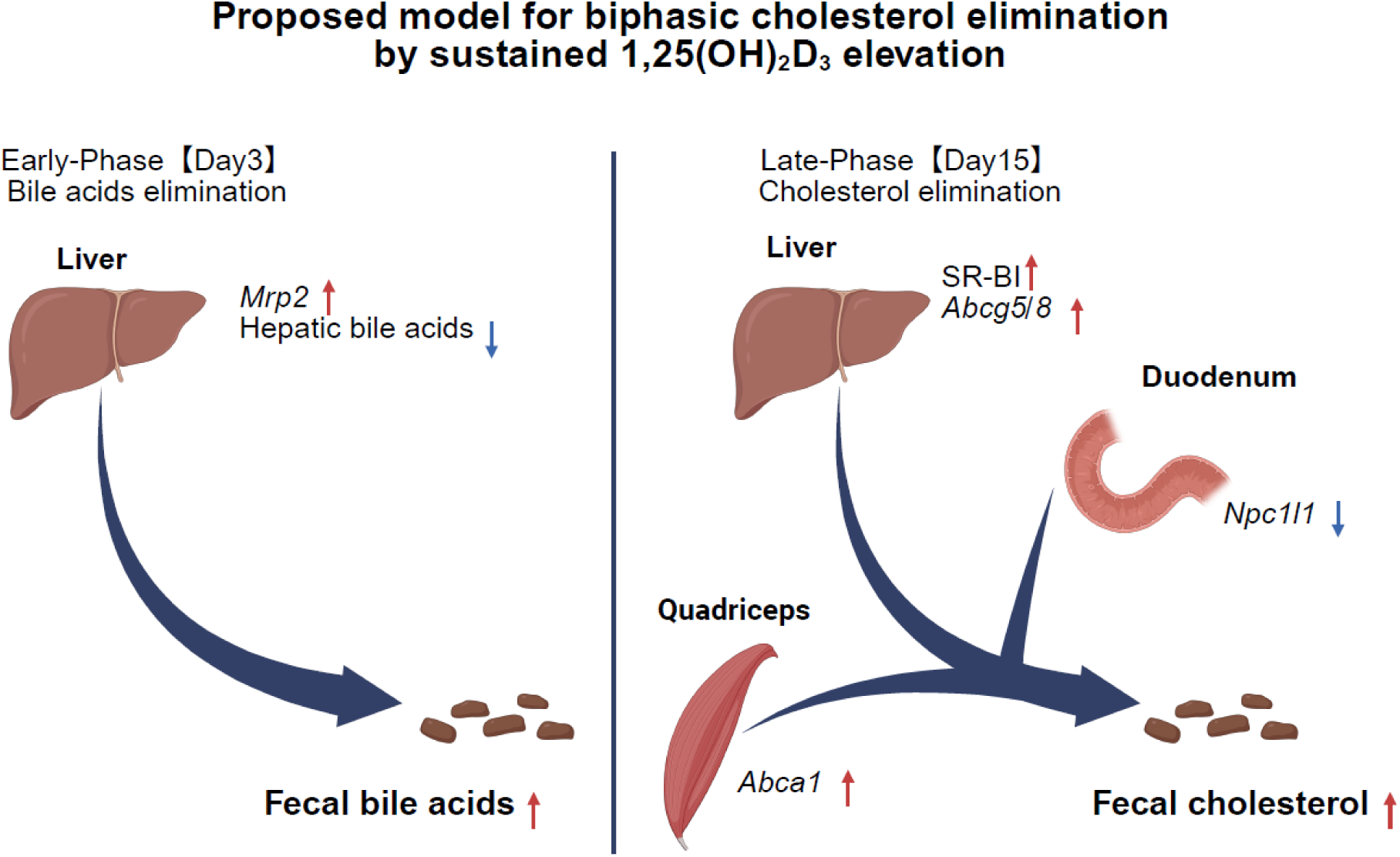
Proposed model for biphasic cholesterol elimination by sustained 1,25(OH)_2_D_3_ elevation. Sustained 1,25(OH)_2_D_3_ elevation promotes biphasic cholesterol elimination, characterized by early fecal bile acid elimination, followed by late fecal cholesterol elimination, along with coordinated changes in hepatic, intestinal, and peripheral cholesterol metabolism. Created in BioRender. Abe, C. (2026) https://BioRender.com/o485dh2

## Author contributions

Hirofumi Sogabe: Methodology, Validation, Formal analysis, Investigation, Writing - Original Draft, Visualization and Data curation.

Chiaki Abe: Conceptualization, Methodology, Validation, Formal analysis, Investigation, Writing - Original Draft, Writing - Review & Editing, Visualization, Data curation, Supervision and Project administration.

Eriko Takaramoto: Validation, Formal analysis and Investigation.

Yo-ichi Nabeshima: Conceptualization, Writing - Review & Editing, Supervision, Project administration, and Funding acquisition.

